# Cannabinoid glycosides: *In vitro* production of a new class of cannabinoids with improved physicochemical properties

**DOI:** 10.1101/104349

**Authors:** Janee’ M. Hardman, Robert T. Brooke, Brandon J. Zipp

## Abstract

The cannabinoid signaling system has recently garnered attention as a therapeutic target for numerous indications, and cannabinoids are now being pursued as new treatment options in diverse medical fields such as neurology, gastroenterology, pain management, and oncology. Cannabinoids are extremely hydrophobic and relatively unstable compounds, and as a result, formulation and delivery options are severely limited. Enzymatic glycosylation is a strategy to alter the physicochemical properties of small molecules, often improving their stability and aqueous solubility, as well as enabling site-specific drug targeting strategies. To determine if cannabinoids are a candidate for glycosylation, a library of glucosyltransferase (UGT) enzymes was screened for glycosylation activity towards various cannabinoids. The UGT76G1 enzyme from *Stevia rebaudiana* has been identified as having glucosyltransferase activity towards a broad range of cannabinoids. Compounds that were successfully glycosylated by UGT76G1 include the phytocannabinoids cannabidiol (CBD), Δ^9^-tetrahydrocannabinol (Δ^9^-THC), cannabidivarin (CBDV), and cannabinol (CBN), and the human endocannabinoids anandamide (AEA), 2-arachidonoyl-glycerol (2AG), 1-arachidonoyl-glycerol (1AG), and synaptamide (DHEA). Interestingly, UGT76G1 is able to transfer primary, secondary, and tertiary glycosylations at each acceptor of most of the cannabinoids tested. Additionally, Os03g0702000p, a glycosyltransferase from *Oryza sativa*, was able to transfer secondary glucose residues onto cannabinoid monoglycosides previously established by UGT76G1. This new class of cannabinoid-glycosides has been termed cannabosides. The compounds have greatly improved solubility in aqueous solutions. This increased aqueous solubility may enable new oral pharmaceutical delivery options for cannabinoids, as well as targeted delivery and release of cannabinoids within the intestines through glycoside prodrug metabolism.

## Introduction

The human endocannabinoid system (ECS) is a broad-spectrum modulator that plays a role in brain plasticity, nociception, inflammation, regulation of stress and emotions, and addiction (Aizpurua-Olaizola, 2017, Hryciw 2016, Gye 2005, Slavic 2013). It includes the endogenous cannabinoids (endocannabinoids), their respective cannabinoid receptors, and the associated enzymes required to synthesize and break down the endocannabinoids (Mackie, 2008). The ECS is a diverse system whose main receptors, CB1 and CB2, are found throughout the body including the central and peripheral nervous systems, the immune system, and gastrointestinal tract (as reviewed recently by Hasenoehrl 2016 and DiPatrizio 2016), the testes, and the heart. Phytocannabinoids from *Cannabis sativa* are capable of modulating ECS receptors and have a long history of medicinal and recreational use (Touw 1981). Δ^9^-Tetrahydrocannabinol (Δ^9^-THC), perhaps the best known psychoactive phytocannabinoid, is only one of over 70 cannabinoids produced by *Cannabis sativa* (Atakan 2012, Welling 2016). Cannabidiol (CBD), a non-psychoactive phytocannabinoid, has recently garnered attention as a potential anti-epileptic, anti-psychotic, and neuroprotectant (Porter 2013, Leo 2016, Zuardi 2012, Iuvone 2009, for review Mechoulam 2002).

Despite their pharmacological promise, cannabinoids are extremely hydrophobic and have very poor solubility in aqueous solutions, necessitating the use of oils and solvents as a carriers in drug formulations, which are not tolerated well and may contribute to oral lesions at high dose or in long-term use (Scully 2007). Numerous efforts have been undertaken to improve the aqueous solubility of CBD and THC. Cyclized maltodextrins have been found to improve the solubility of cannabinoids non-covalently, and covalent modifications have produced chemically derivatized CBD prodrugs with improved solubility (Jarho 1998, WO2009018389, WO2012011112). Fluorine substitutions of CBD have also been created through synthetic chemical manipulations in an effort to functionalize CBD (WO2014108899). Others have gone so far as to modify THC with carboxamido, imidazole, pyrazole, triazole and morpholine pentyl side chain analogs, in addition to converting the phenolic hydroxyl group to substituted esters in an effort to increase solubility (Martin 2006). These strategies have improved the solubility of cannabinoids with limited success, and their unnatural compositions release synthetic prodrug moieties upon hydrolysis.

As with synthetic chemistry, *in vivo* detoxification strategies serve as another model for improving the solubility of cannabinoids. Screening of human liver UGTs against cannabinoids found that cannabinol (CBN) is efficiently glucuronidated by human UGT1A10 *in vitro*, and minor activity was demonstrated towards CBD with UGT1A9 and UGT2B7 (US8,410,064). Screening of plant cell cultures has shown that CBN is glycosylated when incubated with *Pinellia ternata* cell cultures (Tanaka 1993). Similarly, CBD was shown to be glycosylated when incubated with tissue cultures from *Pinellia ternata* and *Datura inoxia*, yielding CBD-6’-O-β-D-glucopyranoside and CBD-(2’,6’)-O-β-D-diglucopyranoside (Tanaka 1996). These cell culture biotransformation studies demonstrated the potential for limited glycosylation of cannabinoids, but previously, no specific enzymes or methods had been identified that could enable the production of high-purity pharmaceutical preparations or a diverse class of cannabinoid-glycosides.

The plant *Stevia rebaudiana* produces a diverse family of steviol glycosides and possesses an untapped pool of UGT enzymes, making Stevia an ideal candidate for small-molecule glycosylation screening. UGT76G1 is a glucosyltransferase from Stevia capable of transferring a secondary glucose to the C3-hydroxyl of the primary glycosylation on both C13-OH and C19-COOH positions of the steviol glycoside, and thus its substrates include steviolmonoside, stevioside, rubusoside, RebA, RebD, RebG, and RebE (Richman et al. 2005, Stevia First Corp unpublished data).

To address the issue of poor cannabinoid solubility and identify methods of producing novel cannabinoid pharmaceutical prodrugs, we screened glucosyltransferase enzymes from Stevia and other organisms to identify candidates for the glycosylation of cannabinoids. We also set out to characterize any resulting cannabinoid glycosides for improved physicochemical properties.

## Results

### UGT76G1 Glycosylation of Cannabidiol

To identify enzymes with glycosylation activity towards cannabinoids, a *Stevia rebaudiana* UGT enzyme library was reacted with CBD (Supplemental figure 1). Upon incubation of CBD with the *Stevia rebaudiana* glucosyltransferase UGT76G1, depletion of the input CBD was observed by RP-HPLC (dotted line, CBD retention time 13.6 min, Figure 1A). CBD was positively identified in the HPLC line trace based on comparison with the retention time of a purchased CBD standard (Cayman Chemical), along with a doublet fingerprint peak at the cannabinoid absorbance maximum of 275 nm. Prolonged incubation of CBD with UGT76G1 yielded 4 distinct glycoside product mobility groups with retention times of 8.75, 9.0, 10.3, and 10.7 minutes, respectively, with other minor products present (Figure 1A). Glycosylation of CBD caused a slight shifting of the fingerprint doublet absorbance maximum at 275 nm to 270 nm, which is consistent with other aglycone-to-glycoside conversions (Figure 1B). CBD glycosylation was found to be UDPG-dependent, and no direct glycosylation of CBD was observed with other tested enzymes. In the presence of excess UDPG, UGT76G1 depleted >95% of the input CBD and had an equilibrium constant (Keq) of ~24. LC-ESI-MS was performed on the CBD glycoside mixture resulting from the UGT76G1 CBD reaction, and [M + H] *m/z* peaks were detected for the CBD aglycone along with the CBD monoside, diglycoside, triglycoside and tetraglycoside (m/z = 315, 477, 639, 801, and 982, respectively, Supplemental Figure 2A). ^1^H-NMR of the purified compounds with HPLC retention times of 9.0 and 10.7 minutes confirmed the production of the CBD diglycosides VB104 and VB110. VB104 is a CBD diglycoside where one available hydroxyl acceptor group has been conjugated with two glucose residues, and its production was confirmed by the presence of two distinct peaks in the aromatic region representing protons 3’ and 5’ from the substituted resorcinol ring of CBD (6.22 and 6.40 ppm, 1H each, Supplemental Figures 2B, 2C). VB110 production was confirmed by the presence of one doublet peak in the aromatic region (6.68 ppm, 2H) representing protons 3’ and 5’, indicating that both the 2’ and 6’ hydroxyl acceptor groups had been conjugated with a glucose residue (Supplemental Figure 2D). The determined CBD glycoside structures of UGT76G1 are depicted in Figure 1D.

**Figure 1:**
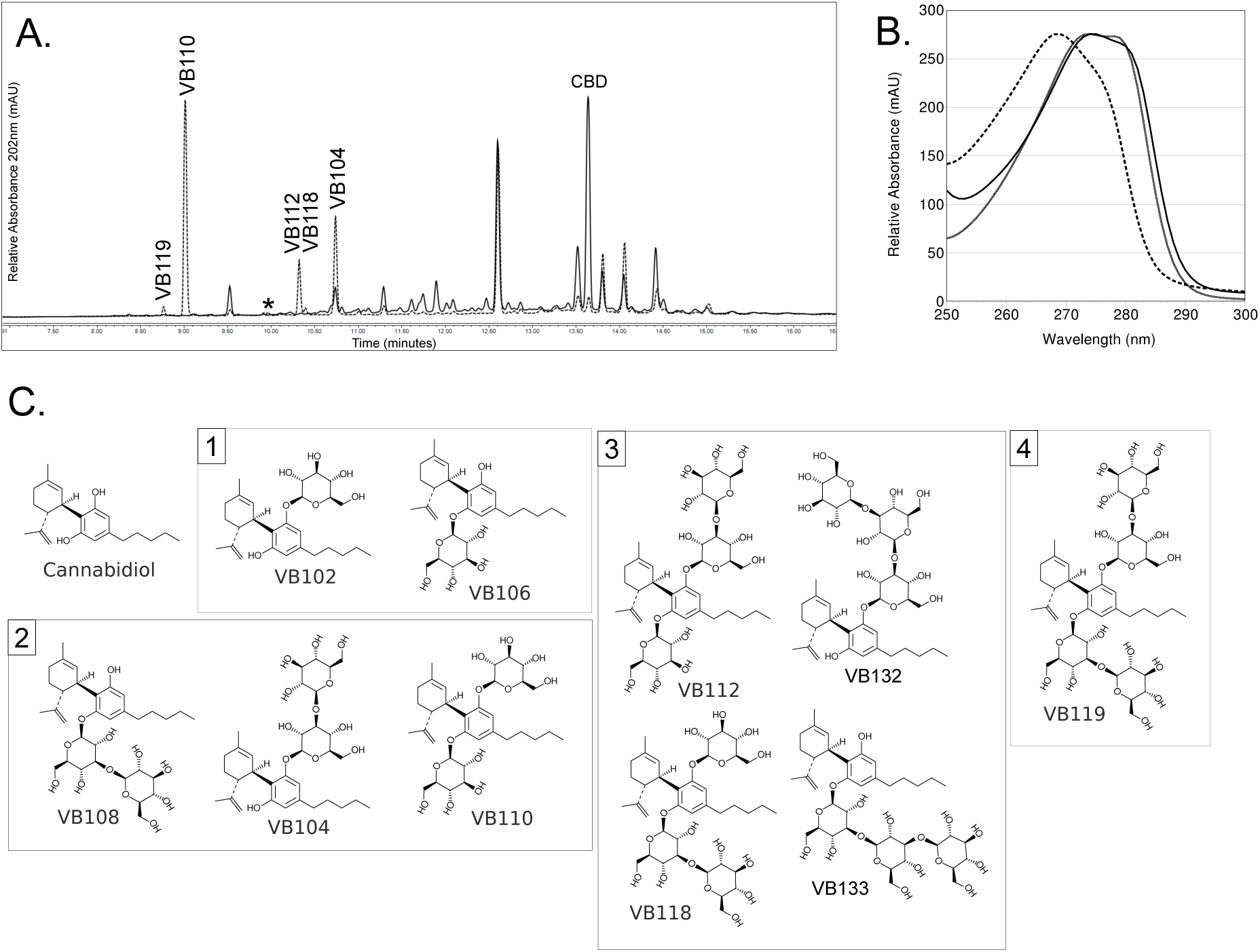
UGT76G1 from *Stevia rebaudiana* glycosylates cannabidiol (CBD) Cannabidiol was reacted with UGT76G1 *in vitro* and the reaction was observed by HPLC (A). The control reaction contained all reagents but lacked only the sugar donor UDPG (solid linetrace). Depletion of CBD was observed upon addition of UDPG, and major product peaks were present at 8.75’, 9.0’, 10.3’, and 10.7’ (dotted linetrace). CBD displayed a broad absorbance doublet centered at 275 nm (solid black line) that is maintained for VB104 (gray line) and shifted for VB110 (dotted line)(B). Structures of CBD and CBD-glycosides as determined by stepwise glycosylation reactions, LCMS-ESI-UV, and H-NMR (C). Product glycosides are organized by the total number of glycosylations present.

### UGT76G1 Glycosylation of Additional Phytocannabinoids

Based on the efficient glycosylation of the hydroxyl groups of CBD by UGT76G1, additional phytocannabinoids were screened as substrates for UGT76G1, including CBDV, CBN, and Δ^9^-THC. CBDV, a cannabinoid differing from CBD by a three-carbon rather than a five-carbon hydrocarbon tail, was selected to be screened next due to its structural similarity to CBD. CBDV was incubated with UGT76G1 and UDPG in a similar manner as CBD. The reaction was monitored by RP-HPLC and CBDV depletion was observed (CDBV retention time 12.7 min), in addition to the appearance of three additional mobility peaks at 8.5, 9.7, and 10.0 minutes (Figure 3A). Formation of these products was dependent on addition of both UGT76G1 and UDPG. The three new products formed displayed the same absorbance characteristics as CBDV and were determined to be the primary glycosides CBDV-2'-O-glucopyranoside or CBDV-6'-O-glucopyranoside, the secondary glycosides CBDV-2'-O-(3-1)-diglucopyranoside or CBDV-6'-O-(3-1)-diglucopyranoside, and CBDV-2’,6’-O-diglucopyranoside (compounds VB202/VB206, VB204/VB208, and VB210 respectively). With additional reaction time it was determined that minor product peaks containing higher-order glycosides were also formed, analogous to what was seen when CBD was the substrate. The reaction proceeded to >95% substrate conversion with K_eq_ ~24.

Both the 2’ and 6’ hydroxyl groups of CBD and CBDV were successfully glycosylated and both compounds exhibit free rotation about the C1’ bond between the resorcinol ring and the terpene ring. To determine if the cannabinoids were binding in the active site of UGT76G1 in a rigid conformation or if there was rotation along the C1’ bond, Δ^9^-tetrahydrocannabinol (Δ^9^-THC) was screened as a potential substrate. Δ^9^-THC was hypothesized to provide insight into active site binding directionality because it has undergone a ring closure between the 6’ hydroxyl group and the terpene ring, thus eliminating any rotation about the C1’ bond. Additionally, Δ^9^-THC only has the 2’ hydroxyl group free for glycosylation (1-OH based on formal dibenzopyran numbering for Δ^9^-THC). Upon incubation of Δ^9^-THC with UGT76G1 and UDPG, RP-HPLC showed depletion of the input Δ^9^-THC (retention time 14.5 min) and the formation of three main product peaks at 10.6, 10.9, and 11.7 minutes (dotted line, Figure 2B). The three new products formed displayed a similar UV-VIS absorbance spectrum to Δ^9^-THC (data not shown). LC-ESI-MS analysis of the glycoside mixture confirmed the identity of the glycosides as Δ^9^-THC monoside, diglycoside and triglycoside ([M + H] *m/*z = 315, 477, 639 and 801, respectively, Supplemental Figure 2D). Structures for Δ9-THC are as depicted in Figure 3D.

**Figure 2:**
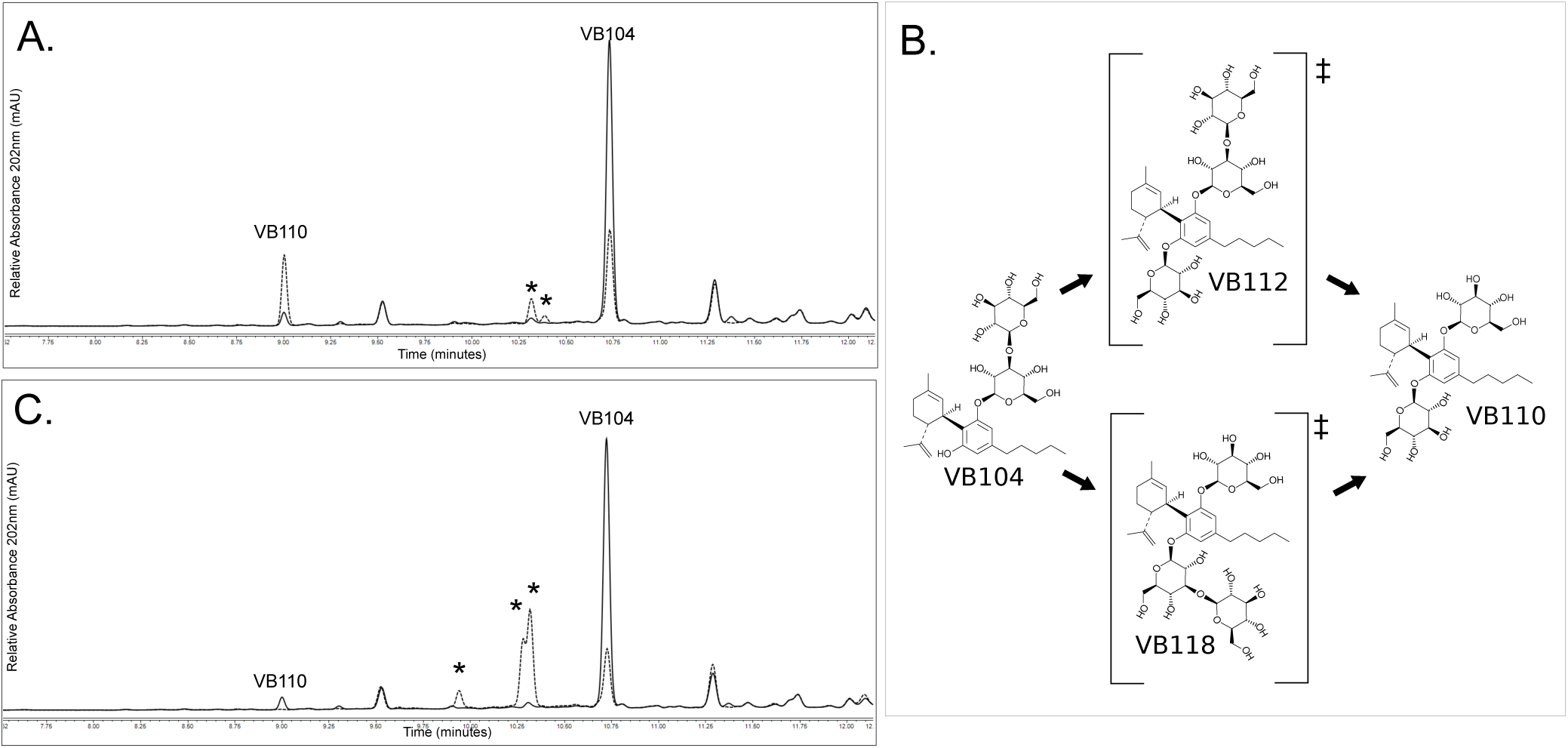
Secondary glycosylation with UGT76G1 and Os03g0702000p. CBD-glycoside VB104 was partially purified and further reacted with UGT76G1 (A). The control reaction contained no UDPG (solid linetrace), with the substrate containing VB104 (10.7’) and a small VB110 peak (9.0’). Upon addition of UDPG, a decrease was seen in VB104 (dotted linetrace), as well as increases in VB110 and two minor peaks at 10.32’ and 10.39’. A proposed biosynthetic pathway from VB104 to VB110 is depicted (B). Additionally, partially purified VB104 was further reacted with Os03g0702000p (C), with inputs as in Figure 2A (solid linetrace). Upon addition of UDPG, VB104 and VB110 decreased in amount, and new peaks were observed at 9.94’, 10.27’, and 10.31’ (dotted linetrace).

**Figure 3:**
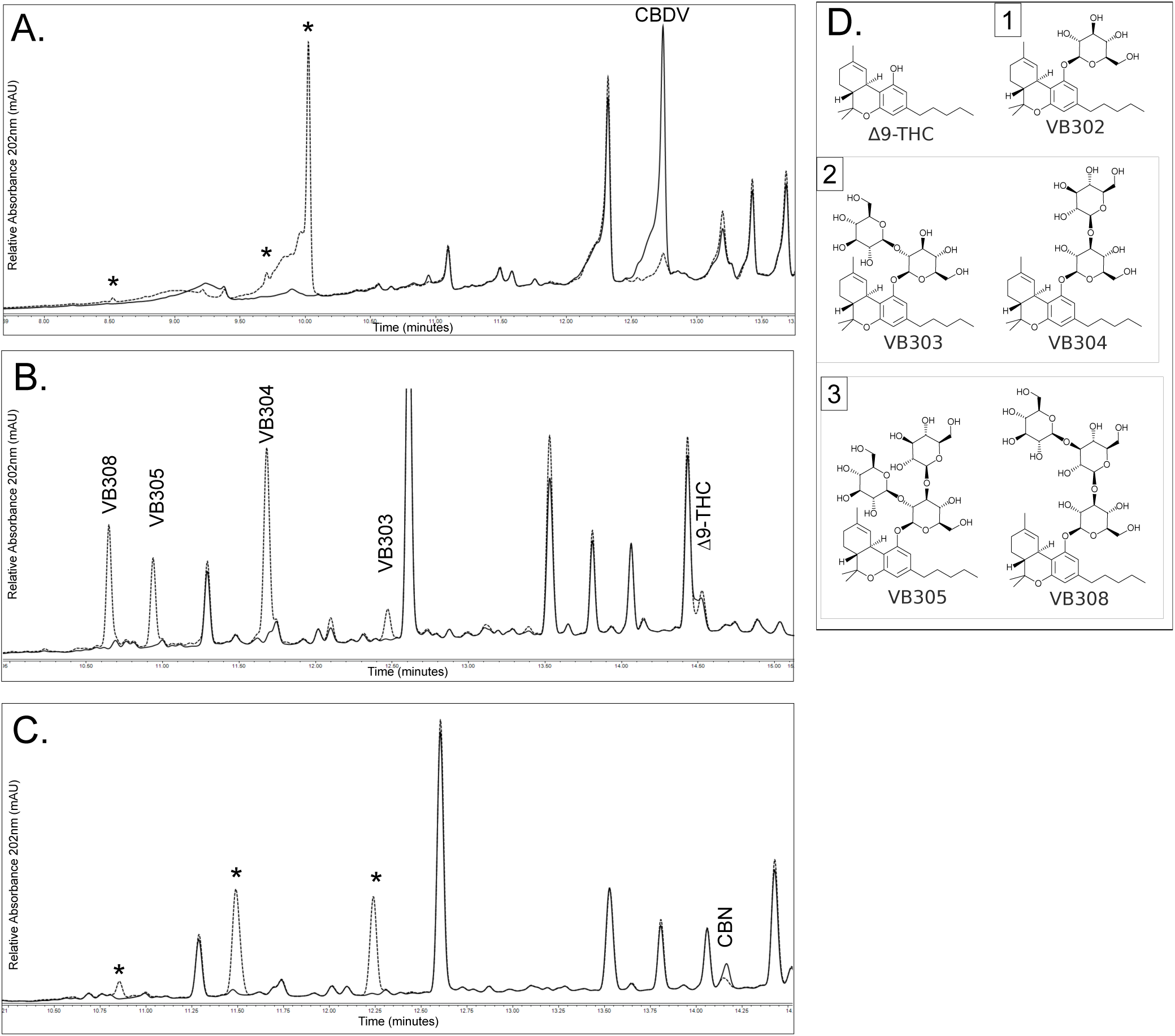
UGT76G1 glycosylates additional phytocannabinoids. Cannabidivarin (CBDV) was reacted with UGT76G1 *in vitro* and the reaction was observed by HPLC (A). The control reaction contained all reagents but lacked only the sugar donor UDPG (solid linetrace). Depletion of CBDV (12.75’) was observed upon addition of UDPG, and major product peaks were present at 8.52’, 9.7’, and 10.0’ (dotted linetrace). Δ9-Tetrahydrocannabinol (THC) was reacted with UGT76G1 *in vitro* and the reaction was observed by HPLC (B). The control reaction contained all reagents but lacked only the sugar donor UDPG (solid linetrace). Depletion of THC (14.52’) was observed upon addition of UDPG, and major product peaks were present at 10.64’, 10.93’, 11.68’, and 12.48’ (dotted linetrace). Cannabinol (CBN) was reacted with UGT76G1 *in vitro* and the reaction was observed by HPLC (C). The control reaction contained all reagents but lacked only the sugar donor UDPG (solid linetrace). Depletion of CBN (14.17’) was observed upon addition of UDPG, and major product peaks were present at 10.85’, 11.48’, and 12.23’ (dotted linetrace). Structures of THC and THC-glycosides as determined by stepwise glycosylation reactions, and LCMS-ESI-UV (D). Product glycosides are organized by the total number of glycosylations present.

The final phytocannabinoid that was screened as a substrate for UGT76G1 was the Δ^9^-THC natural degradation product, cannabinol (CBN), which resembles Δ^9^-THC but the upper terpene ring has been fully oxidized to its aromatic form. Upon incubation of CBN with UGT76G1 and UDPG, RP-HPLC showed depletion of the input CBN (CBN retention time 14.2 min) and formation of three main product peaks at 10.8, 11.5 and 12.2 minutes (Figure 3C). The three new products displayed the same absorbance characteristics as CBN (data not shown).

### Secondary Glycosylation With Os03g0702000p

Branched chain glycosides are often glycosylated by specific enzymes in a stepwise manner, and certain UGTs only recognize a glycoside as their substrate. As UGT76G1 was found to establish primary, secondary, and tertiary glycosylations on the hydroxyl groups of CBD, additional UGTs were screened for their ability to recognize cannabinoid glycosides as substrates for further glycosylation. Os03g0702000p (formerly referred to as EUGT11) is a UGT from *Oryza sativa* that has secondary glycosylation activity towards steviol glycosides (WO 2013022989). It was tested and determined that Os03g0702000p is capable of transferring an additional glucose moiety from UDPG onto the C2-hydroxyl of the primary sugar established by UGT76G1 (β-2→1 connectivity, Figure 2C), similar to the secondary glycosylation activity that UGT76G1 has towards the C3-hydroxyl of the primary glucose residue glycosylation (β-3→1 connectivity). Upon incubation of a CBD glycoside mixture generated by UGT76G1 with Os03g0702000p, four additional glycoside products were formed with HPLC retention times of 9.9, 10.2, 10.3, and 10.6 minutes, which retained the cannabinoid absorbance maxima at 270 nm. This glycosylation activity is consistent with the activity of UGT Os03g0702000p towards steviol glycosides in establishing C2-hydroxyl secondary glycosylations (β-2→1 connectivity) on existing primary glucose residues. Similar glycosylation activity was observed when CBDV, Δ^9^-THC, and CBN glycoside mixtures were incubated with Os03g07000p and excess UDPG (data not shown). All glycoside products were identified based on the retained cannabinoid absorbance maxima at 275 nm. No glycosylation activity was seen when Os03g0702000p alone was incubated with cannabinoids.

### UGT76G1 Glycosylation of Endocannabinoids

As UGT76G1 has been determined to recognize a broad class of phytocannabinoids, it was hypothesized that the same enzyme active site may also accommodate and glycosylate endocannabinoids, the endogenous human signaling molecules recognized by the cannabinoid receptors CB1 and CB2 in humans. Accordingly, the endocannabinoids AEA, 2AG, 1AG, and DHEA were obtained and screened as possible substrates for glycosylation by UGT76G1. All endocannabinoid compounds showed UDPG-dependent depletion of the substrates and the formation of new product peaks by RP-HPLC (Figures 4A, 4B, and 4C, respectively). Upon incubation with UGT76G1 and UDPG, AEA (retention time of 13.9 min) yielded one glycoside product peak with a retention time of 12.5 min (Figure 4A). Glycosylation of AEA occurred slowly, and reactions did not proceed to product/substrate ratios seen with phytocannabinoids. 2-AG and 1-AG were obtained in a 90:10 2-AG:1-AG ratio and each displayed an absorbance maximum at 233 nm and RP-HPLC retention times of 14.2 (1-AG) and 14.3 (2-AG) minutes. Upon incubation with UGT76G1 and UDPG, both compounds showed UDPG-dependent depletion by UGT76G1 (Figure 4B). Nine new arachidonoylglycerol-glycoside product peaks were observed on the HPLC trace at 11.4, 11.7, 11.8, 11.9, 12.0, 12.5, 12.9, 13.1, and 13.3 minutes, and all retained the signature 233 nm absorbance maxima (Figure 4B). Finally, DHEA, which has an RP-HPLC retention time of 13.7 min and an absorbance maximum of 237 nm, was incubated with UGT76G1 and UDPG. Two new glycoside product peaks were observed in the HPLC line trace at 10.1 and 12.4 minutes, and were identified as glycosides of DHEA as both displayed the signature absorbance maximum of 237 nm (Figure 4C).

**Figure 4:**
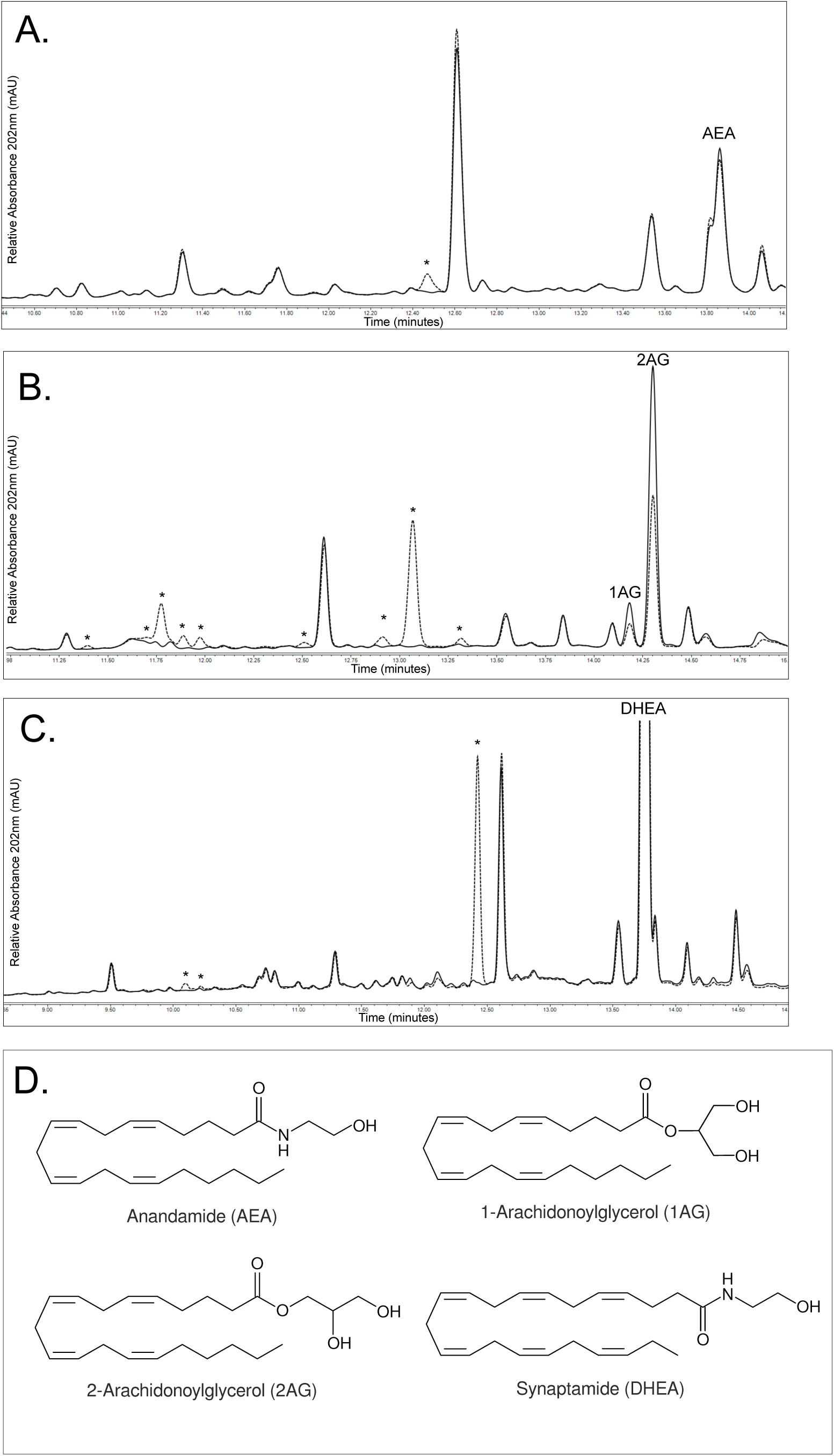
UGT76G1 glycosylates endocannabinoids. Anandamide (AEA) was reacted with UGT76G1 *in vitro* and the reaction was observed by HPLC (A). The control reaction contained all reagents but lacked only the sugar donor UDPG (solid linetrace). Depletion of AEA (13.85’) was observed upon addition of UDPG, and the major product peak was present at 12.47’ (dotted linetrace). A 9:1 mixture of 2-arachidonoyl-glycerol (2AG) and 1-arachidonoyl-glycerol (1AG) was reacted with UGT76G1 *in vitro* and the reaction was observed by HPLC (B). The control reaction contained all reagents but lacked only the sugar donor UDPG (solid linetrace). Depletion of 2AG (14.31’) and 1AG (14.18’) was observed upon addition of UDPG and major product peaks were present at 11.39’, 11.70’, 11.77’, 11.89’, 11.97’, 12.51’, 12.92’, 13.07’, and 13.32’ (dotted linetrace). Synaptamide (DHEA) was reacted with UGT76G1 *in vitro* and the reaction was observed by HPLC (C). The control reaction contained all reagents but lacked only the sugar donor UDPG (solid linetrace). Depletion of DHEA (13.75’) was observed upon addition of UDPG, and the major product peak was present at 12.42’ with minor peaks at 10.09’, and 10.21’ (dotted linetrace). (D). Endocannabinoid substrates of UGT76G1.

### Physicochemical properties of cannabinoid glycosides

Phytocannabinoids are hydrophobic compounds, which limits their formulation and delivery options for pharmaceutical uses. The cannabinoid glycoside products in the present study all displayed advanced elution from the reverse-phase HPLC separation, suggesting that they are less hydrophobic than their precursors. Based on this observation, it was hypothesized that the cannabinoid glycoside products would have improved aqueous solubility. The standard quantification for chemical solubility is partitioning in water and octanol
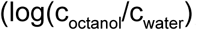, or ClogP), and recently software has been developed to estimate the ClogP values. Select cannabosides were analyzed and ClogP values were calculated (Table 1). The addition of glucose residues to the cannabinoid backbone significantly decreased the ClogP relative to the parent compounds. Δ^9^-THC had a ClogP value of 7.2, but the addition of one glucose residue (VB302) decreased the CLogP value to 5.7, and the addition of two glucose residues (VB304) reduced the value even further to 4.7. To explore how the location of the added sugar residues also affects the CLogP value, CLogP values for CBD were calculated. Compared to Δ^9^-THC, which has one hydroxyl acceptor group, CBD has two hydroxyl acceptor groups at C2’ and C6’. The calculated CLogP value for CBD was 6.6, and the addition of one glucose residue reduced this value to 5.1. With addition of a diglycoside to the hydroxyl group C2’ (VB104), the CLogP value decreased even further to 4.3. When one glucose residue was added to each of the C2’ and C6’ hydroxyl groups (VB110), the CLogP value decreased to 3.4. The ClogP values predicted *in silico* were comparable to values determined experimentally by RP-HPLC using compounds with known ClogP values as standards. Comparison of the predicted ClogP values for the cannabosides with their elution times revealed a linear relationship between the ClogP value and the advanced elution time (Figure 5A). The decreases in CLogP values observed for the cannabinoid glycosides indicate improved water solubility of cannabosides compared to the parent aglycones.

**Figure 5:**
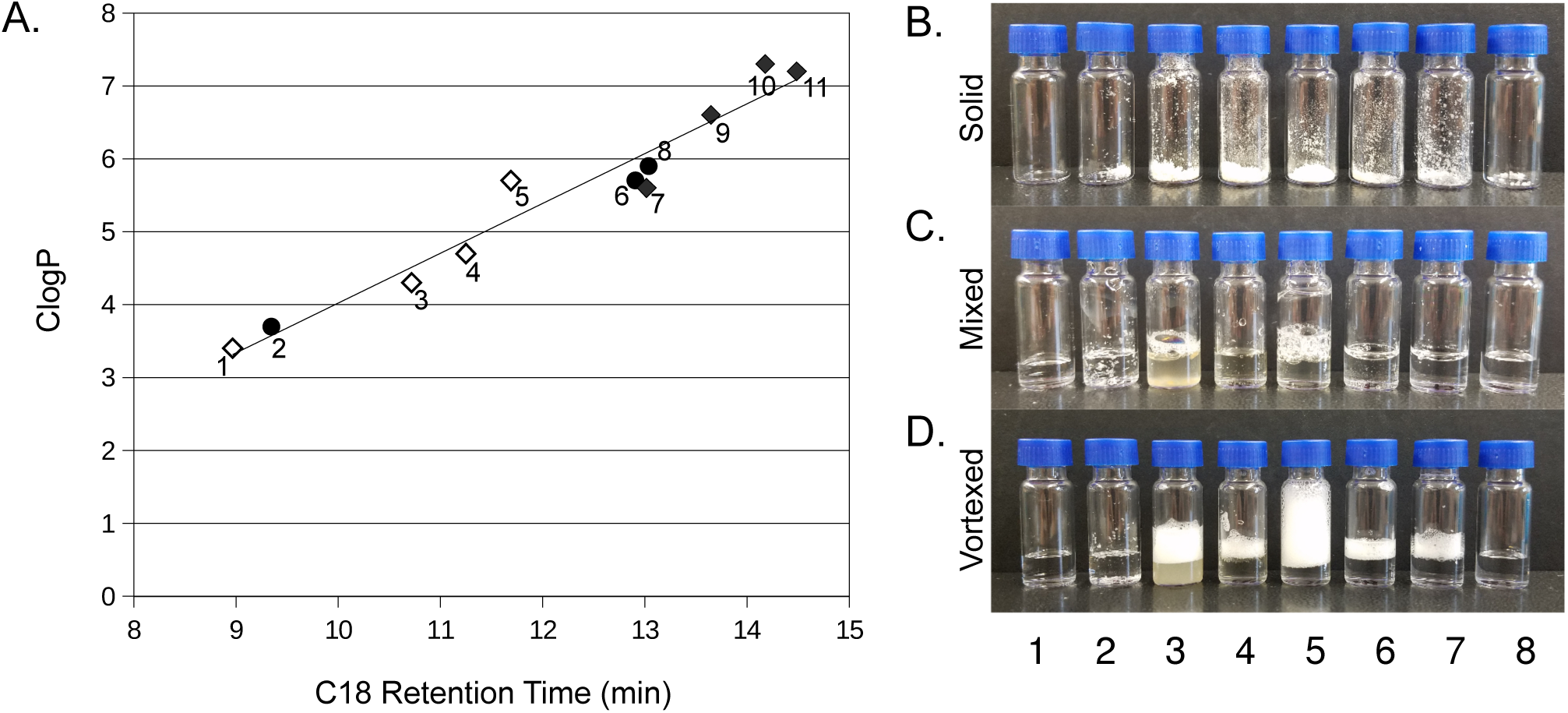
Physicochemical properties of cannabosides. ClogP values for select cannabosides, cannabinoids, and cannabinoid-metabolites from Table 1 were plotted against RP-HPLC retention times to assess linearity (A). The C18 retention times for cannabinoids (filled diamonds), cannabosides (open-diamonds), and cannabinoid-metabolites (filled circles) were plotted against ClogP values from Table 1 (A). Linear regression was performed on all data points (R2 = 0.9455) and plotted as the line in (A). Aqueous solubility and detersive properties are displayed (B, C, and D). Solids in vials (B), hydrated and mixed by pipetting at 1% in water (C), and following 1 minute of vortexing (D). Vials are as follows: 1. Water, 2. CBD, 3. VB104, 4. VB110, 5. VB304, 6, Saponin, 7. SDS, and 8. Sucrose.

**Table 1:** Physicochemical properties of cannabosides, cannabinoids, and cannabinoid-metabolites. The RP-HPLC retention times were determined empirically, and the ClogP values were calculated in silico for a subset of cannabosides, cannabinoids, and cannabinoid-metabolites.

| # | Compound | ClogP |
| --- | --- | --- |
| 1 | VB110 | 3.4 |
| 2 | 11-COOH-Tetrahydrocannabinol Glucuronide | 3.7 |
| 3 | VB104 | 4.3 |
| 4 | VB304 | 4.7 |
| 5 | VB302 | 5.7 |
| 6 | 11-COOH-Tetrahydrocannabinol | 5.7 |
| 7 | Cannabidivarin | 5.6 |
| 8 | 11-OH-Tetrahydrocannabinol | 5.9 |
| 9 | Cannabidiol | 6.6 |
| 10 | Cannabinol | 7.3 |
| 11 | Tetrahydrocannabinol | 7.2 |

Because the ClogP values and C18 retention times indicated that cannabosides have decreased hydrophobicity, the aqueous solubility was directly tested (Figure 5B, C). Solids in vials were imaged after hydration and mixing by pipetting at 1% in water, followed by 1 minute of vortexing (Figures 5B, C, and D, respectively). The vials contained: 1. water (for reference), 2. CBD (negative solubility control), 3. VB104, 4. VB110, 5. VB304, 6, saponin (positive foaming control), 7. SDS (positive foaming control), and 8. sucrose (positive solubility, negative foaming control). Pure crystalline CBD was insoluble and water failed to solubilize it. Cannaboside VB104 was hydrated by the added water but the solutions remained cloudy and opaque; additionally, it possessed detersive properties (Figure 5D vial 3). VB110 was fully solubilized in water and created a translucent dark liquid. VB304 had partial solubility in water and displayed significant detersive properties. Saponins and SDS were both water soluble and detersive at 1% in water. Finally, sucrose was fully water soluble at 1% and did not display any foaming. The significant foaming shown by VB304 persisted for over 8 hours, whereas the SDS foaming subsided after 1 hour (data not shown).

## Discussion

In the present study, UGT76G1 was capable of transferring primary, secondary, and tertiary (β 3-1) glycosylations onto receptor molecules including phyto- and endocannabinoids. It was also found that UGT76G1 is able to specifically glycosylate both the C2' and C6' hydroxyl groups on CBD and CBDV, in addition to the C1 hydroxyl group of Δ^9^-THC and CBN. Os03g0702000p from *Oryza sativa* was capable of transferring secondary (β 2-1) glycosylations onto primary glucose residues established by UGT76G1. The cannabinoid-glycosides produced by these reactions display greatly improved solubility relative to the aglycone precursors and represent a new class of water-soluble cannabinoids termed cannabosides.

The reactions for production of cannabosides are dependent on UDPG, and the products maintain a cannabinoid absorbance spectrum. Multiple CBD-glycoside product peaks are seen on HPLC, and stepwise glycosylation and structural characterization have shown that the products consist of CBD mono, di-, tri-, and tetra-glycosides. All of the CBD glycosides show advanced elution from the C18 column relative to the CBD substrate, indicating a decrease in hydrophobicity. Calculation of the ClogP values for the predominant CBD-glycosides show greatly decreased hydrophobicity compared to the CBD aglycone.

The number of CBD-glycoside product mobility groups indicates multiple glycosylations that occur on both hydroxyl groups. To determine how cannabinoids are coordinated in the active site of UGT76G1, CBD was superpositioned over the bi-functional substrate for UGT76G1, rebaudioside E (RebE) (Supplemental Figure 3). The mechanism for dual hydroxyl glycosylation is likely one of two possibilities: either CBD is docking in the UGT76G1 active site both forwards and backwards, creating a cis-like sugar conformation for the glycosylations relative to the cannabinoid backbone (mechanism depicted in Supplemental Figure 4), or alternatively, the rotational freedom about the bond at C1' may allow the resorcinol-hydroxyl-glycoside group to rotate 180 degrees after glycosylation, placing the second free hydroxyl group in the active site for glycosylation that occurs in a trans-like conformation relative to the first glycosylation (C6 carbon as described by Mazur 2009, mechanism depicted in Supplemental Figure 5). The difference between these two mechanisms is slight, but they produce structurally distinct glycoside products that may have significantly different properties.

The majority of reactions with UGT76G1 and CBD yielded only diglycosides and higher. Tanaka observed CBD-glycosides after incubating CBD in root cell cultures of *Pinellia ternata* for periods up to 30 days and did not perform any short-term assays (Tanaka 1996). As such, Tanaka et al. described difficulty in identifying monoglycosylations of cannabidiol in their biotransformation reactions. Similarly, in this work, UGT76G1 did not produce single glycosylations of CBD in prolonged *in vitro* enzymatic reactions. This is likely due to the high affinity of CBD-monoglycosides towards the active site of UGT76G1. Short-term kinetic assays with UGT76G1 did produce CBD-monoglycosylations (VB102, VB103) when reactions were stopped using 80% acetonitrile (data not shown). CBD may require two glycosylations to achieve adequate aqueous solubility for escaping the active site of UGT76G1.

Performing enzyme kinetics with UGT76G1 and CBD was complicated by the unique biochemistry of UGT76G1. First, the cannabinoid glycoside products are also substrates for further glycosylation by UGT76G1; therefore, the reactions produce multiple competitive inhibitors very early during the reaction. It was also observed that excess CBD acted to substantially inhibit UGT76G1 reactions; therefore, the substrate concentration needed to be limited to prevent kinetic inhibition of the enzyme (data not shown). Based on this observation, a fed-batch conversion reaction in which CBD was dosed in small amounts on an hourly basis was performed to determine whether maintaining a low concentration of CBD was beneficial to the conversion reaction. Interestingly, this fed-batch reaction preferentially produced VB104, indicating that CBD is the preferred substrate for UGT76G1 when competing with VB104. It was also observed that pre-incubation of UGT76G1 with UDPG resulted in a burst phase of glycoside production that quickly leveled off and appeared to reach a linear, steady-state rate of production of VB104. This initial burst phase likely exhausted the pre-bound UDPG, after which the dissociation of UDP from the active site and concurrent binding of UDPG came to equilibrium.

UGT76G1 exhibited glycosylation activity towards THC that was similar to its activity towards CBD. The multiple THC-glycoside products showed advanced elution from the C18 column and maintained the cannabinoid absorbance spectrum. Similar to CBD, the primary products of UGT76G1 glycosylation were mostly THC-diglycosides and higher. Because LC-ESI-MS clearly revealed a THC-triglycoside, UGT76G1 is capable of transferring up to tertiary glycosides to the same recipient substrate. The fourth product peak was determined to be a glycoside, but the precise structure was not elucidated at the time this manuscript was prepared. It is unlikely to be a THC-tetraglycoside, as the LC-ESI-MS did not indicate a tetraglycoside molecular weight; rather, it may be a secondary glycosylation other than (β 3-1) attachment, or a degradation product such as CBN-glycoside. Given that the rigid structure of Δ^9^-THC does not have the same rotational freedom as CBD around the C1' resorcinol ring attachment, the cannabinoid backbone is recognized in the active site of UGT76G1 with the Δ^9^-THC C1 hydroxyl group situated towards the UDPG sugar donor (pyran numbering, Figure 1B).

As originally hypothesized, cannabosides have greatly improved solubility in aqueous solutions. The solubility is dependent on the number of sugars attached, as well as the position and attachment site of the individual sugars. The highest solubility was observed for VB119 and VB110, CBD-glycosides with glucose molecules on opposing C2’ and C6’ hydroxyl groups. For VB110, it was found that solutions in excess of 50% w/v (0.78 M) were possible using only water as the solvent (data not shown). Factoring in the molecular mass addition resulting from the two attached sugars, the CBD content is 24.6% by mass, or 0.39 M. This empirically determined aqueous solubility is reinforced by ClogP values calculated *in silico* using ChemDraw Ultra for phytocannabinoids conjugated with a maximum of two glucose residues. The ClogP values also show that the number and position of the glycosylations impact the predicted solubility, and validate the aqueous solubility differences seen between the two CBD-diglycosides VB110 (ClogP = 3.4) and VB104 (ClogP = 4.3). Whereas VB110 has glycosylations on opposite sides of the molecule, VB104 is more amphipathic with a hydrophilic glycosylation opposed by the free hydroxyl group. This is more analogous to the THC-glycosides that have lost the free hydroxyl group to ring closure and exhibit a more severely amphipathic structure. During the preparation of aqueous solutions, it was observed that specific cannabosides displayed detersive properties as indicated by foaming while mixing. This detersive foaming is reminiscent of saponins, which are naturally occurring amphipathic plant glycosides known for their detersive properties, as well as foaming displayed by steviol glycosides (unpublished data). The THC-glycoside VB304 was the most significant detersive molecule, showing greatly increased foaming over saponins and SDS at the same concentration (Figure 5D). Interestingly, this foaming was more stable than that of other cannabosides, saponin, and SDS, and persisted for more than 8 hours.

Phytocannabinoids are known to act as agonists, antagonists, and inverse agonists of the human cannabinoid receptors, disrupting the endogenous cannabinoid signaling. This competition at the same active site indicates a functional and structural similarity between phytocannabinoids and endocannabinoids. Based on these similarities, the most well-characterized endocannabinoids were tested, and UGT76G1 was shown to glycosylate all endocannabinoids evaluated. A clear preference was observed for 2-AG and 1-AG compared with the ethanolamides AEA and DHEA. The weak glycosylation activity of UGT76G1 towards anandamide may also be due to non-optimal reaction conditions such as pH, temperature, or specific buffer chemistry, or the ester group of the arachidonoylglycerols may be preferred over the amide group adjacent to the acceptor hydroxyl group of the ethanolamides. Further experimentation with additional endocannabinoids may shed light on the molecular components that contribute to binding within the UGT76G1 active site.

CBD has been published to degrade to THC, CBN, and quinone derivatives by light, heat, and acidic or basic conditions (for review Mechoulam and Hanus, 2002). Further studies have shown that orally administered CBD may degrade into the psychoactive THC in simulated gastric fluids containing 1% SDS (Watanabe 2007, Merrick 2016). More recently, these *in vitro* results have been called into question as there is no *in vivo* evidence to support these claims (Russo 2017, Vitality Bio unpublished results). In the event that CBD is hydrolyzed to the psychoactive THC by gastric fluids, conjugating sugars to the free hydroxyl groups may protect cannabinoids from conditions that might otherwise produce unwanted cannabinoid byproducts within the stomach.

A growing body of evidence shows that glycosides are capable of acting as prodrugs and have direct therapeutic effects. One example of a class of glycoside pharmaceutical prodrugs is senna glycosides (Senokot, Ex-Lax), which are administered orally, followed by decoupling of the sugars in the large intestine by β-glucosidases secreted from intestinal microbiota (Hardcastle 1970). Site-specific delivery of steroid glycosides to the colon has also been demonstrated (Friend 1985, Friend 1984). Glycosylation of steroids enabled survival of stable bioactive molecules in the acidic stomach environment and delivery into the large intestine, where the aglycones were decoupled by glycosidases and absorbed into the systemic circulation. Colon-specific delivery of cannabosides and decoupling by local β-glucosidases may enable treatment of colon-specific disorders such as inflammatory bowel disease (IBD), including Crohn’s disease and ulcerative colitis (Kunos 2004). In addition to facile delivery to the colon, β-glycosidases are also present universally in different tissues; therefore, delivery of cannabosides by methods that bypass the digestive tract and colon, such as intravenous or intranasal delivery, may enable delivery of cannabinoid aglycones to cells and tissues with adequate expression of glucosidases (Conchie 1959). Additionally, increasing the diversity and complexity of sugar attachments of cannabosides may provide altered distribution and pharmacokinetics *in vivo*, such as for extended- or delayed-release applications depending on the kinetics of glycoside decoupling. Glycoside prodrugs may thus enable site-specific and tissue-specific drug delivery, which would release simple glucose sugars upon prodrug decoupling. Cannabosides used as prodrugs could ultimately enable novel methods of delivering cannabinoids within the body, including the potential to limit the entry of psychoactive compounds into the bloodstream or brain through well-tolerated oral drug formulation and targeted delivery.

In the course of the present work, it was discovered that UGT76G1 from *Stevia rebaudiana* is capable of glycosylating a diverse range of substrates beyond steviol glycosides, including both aglycones and glycosides of multiple cannabinoid species. This promiscuous activity is in agreement with data published during the preparation of this manuscript showing that UGT76G1 may be active towards a wide range of substrates ranging from small aglycones to larger glycosides (Dewitte 2016).

In summary, this collection of novel cannabosides represents a greatly expanded class of cannabinoids with improved physicochemical properties and a wide variety of potential therapeutic applications. Further studies are currently underway to determine the safety and efficacy of these compounds for use as cannabinoid prodrugs.

## Materials and Methods

### General Materials and Methods

Reverse-phase high-performance liquid chromatography (RP-HPLC) was performed on a Dionex 3000 LC system using a Phenomenex Kinetex 5µm XB-C18 100 Å, 150 x 4.6 mm column. Conditions were as follows: column temperature 30°C, autosampler 5°C, flow rate 1.0 mL/min, running time 25 min, solvent A acetonitrile, solvent B water. Solvent A was started at 10% for 4’, then ramped to 99% over 10’, held at 99% for 4’, ramped down to 10% over 2’, and held at 10% for 5’. Detection at 202 nm provided the best information for all cannabinoids and cannabinoid glycosides. Cannabinoids and cannabinoid glycosides were identified using reference standards and the observation of a doublet peak at the cannabinoid absorbance maximum of 275 nm. ^1^H-NMR spectroscopy was performed on a Bruker Avance II 400 MHz instrument (Emery Pharma) with CD_3_OD as the solvent, using TopSpin acquisition and processing software. Liquid chromatography-electrospray ionization mass spectrometry (LC-ESI-MS) analysis was conducted on a Shimadzu LCMS 2010EV instrument (Emery Pharma). LC separation was performed using a Silia Chrom XDB C18 5um, 150A, 4.6X50 mm. The method was 12 minutes with a 5 to 95 water:acetonitrile gradient elution. Low-resolution MS was performed in both positive and negative modes. Acetic acid and formic acid were used as sample additives during analysis, and the injection volumes ranged from 5 to 20 μl.

### Cloning, Expression, Purification, and Characterization of UGT76G1 and Os03g0702000p

A *Stevia rebaudiana* transcriptome library was created. mRNA was extracted from mature stevia leaves using a Spectrum Plant Total RNA extraction kit (Sigma Aldrich). Library preparation and next-gen sequencing were performed on an Illumina HiSeq2000 (Genewiz). Sequence reads were assembled using CLC Genomics Workbench 5.5.1 (Genewiz). The resulting transcriptome was queried with known Stevia UGT proteins via TblastN searches on a local BLAST server. The cDNA of UGT76G1 was amplified from mRNA via PCR utilizing the Bio-Rad *i*Script cDNA Synthesis Kit. Os03g0702000p was synthesized (IDT DNA) and PCR to amplify the genes of UGT76G1 and Os03g0702000p was performed utilizing primers listed in Supplementary S1. The primers were designed to add a 3’ HQ tag to UGT76G1 and a 5’ HQ tag to Os03g0702000p. Reaction mixtures included cDNA (2 μl), forward and reverse primers (10 μM each), dNTPs (200 μM), Phusion HF buffer (1X), and Phusion DNA Polymerase (0.5 μl, 1.0 unit). The amplified sequences were inserted into the inducible pLATE11 vector (Thermo Scientific) using aLICator Ligation Independent Cloning and Expression System Kit 1. The resulting plasmids were used to transform E. coli BL21(DE3) competent cells (New England BioLabs). Positive transformants for each enzyme were confirmed by colony PCR using LIC forward and reverse primers.

Transformant cells for each enzyme were cultivated in 2xYT growth medium containing 100 μg/ml ampicillin while shaking at 20°C overnight. Then, 1.6 ml of inoculum was added to flasks containing 250 ml 2xYT growth medium with 100 μg/ml ampicillin. Cultures were grown to an optical density of 0.6 at 595 nm. UGT76G1 and Os03g0702000p expression was induced by addition of isopropyl-1-*b*-D-thiogalactoside (IPTG) to a final concentration of 1 mM, followed by cultivation at 20°C while shaking for an additional 24 hours.

The cells were harvested by centrifugation at 8,000g for 15 min, then resuspended in Bead Wash Buffer containing 50 mM potassium phosphate (pH 8.0) and 300 mM sodium chloride (NaCl). The cells were lysed by 2 passes through a microfluidizer, and the cellular debris was then removed by centrifugation. The supernatants were applied to Ni-NTA affinity columns (McLab) that had been equilibrated with Bead Wash Buffer. The columns were washed with 10 bed volumes of Wash Buffer containing 50 mM potassium phosphate (pH 8.0), 300 mM NaCl, and 2 mM imidazole. The 3’ HQ tagged UGT76G1 and 5’ HQ tagged Os03g0702000p were eluted from the Ni-NTA columns with 10 bed volumes of Elution Buffer containing 50 mM potassium phosphate (pH 8.0), 300 mM NaCl, and 250 mM imidazole. Sterile glycerol was added to each enzyme eluate to make a 50% UGT76G1 and Os03g0702000p glycerol stock solution, which was stored at −20°C. Characterization and purity analysis of the fractions eluted with 250 mM imidazole were performed by sodium dodecyl sulfate (SDS)-polyacrylamide gel electrophoresis (PAGE) using a 10% gel. Protein concentrations were determined by Bradford assay.

### General Procedure for Glycosylation of Phytocannabinoids and Endocannabinoids with UGT76G1

Glycosylation of selected phytocannabinoids and endocannabinoids by UGT76G1 was performed on a 200 μl scale as proof of concept. The reactions were performed as follows. 1 μg of the selected phytocannabinoid or endocannabinoid substrate (1 μl of a 1 mg/ml solution dissolved in methanol, 0.016 mM final concentration) was added to 200 μl of a reaction mixture containing 50 mM potassium phosphate buffer (pH 7.2), 3 mM magnesium chloride (MgCl_2_), 2.5 mM uridine diphosphate glucose (UDPG) as the sugar donor, and 10 ml of a 50% UGT76G1 glycerol stock solution (1.5 mg/ml stock solution, 1.44 μM final protein concentration). The reaction mixtures were incubated at 28 °C while shaking at 180 rpm for 18 hrs. To stop the reaction, the reaction mixture was extracted three times with 200 μl of ethyl acetate each time. The organic layers containing cannabinoid glycosides and any unreacted substrate were combined, and the ethyl acetate was removed under vacuum; then, the samples were redissolved in 100 μl of 50% MeOH by volume. Reaction products were analyzed by RP-HPLC on a Dionex 3000 LC system using a Phenomenex Kinetex 5 μm XB-C18 100Å column (150 x 4.6 mm). The conditions were as follows: injection volume of 50 μl; column temperature 30 °C; autosampler 8 °C; flow 1 ml/min; running time 25 min; solvent A deionized water (di-H_2_O), solvent B acetonitrile (ACN), gradient elution 10-99% ACN.

Glycosylation reactions for the phytocannabinoids CBD and THC were scaled up proportionally to 100 ml to obtain an adequate amount of glycosylated products for characterization by LC-MS and ^1^H NMR. UGT76G1 was deactivated and precipitated from the glycosylation reaction mixture by treatment at 95 °C for 10 minutes. Clarified reaction mixture was obtained by centrifugation. Purification and separation of the glycosylated products is described below.

### Glycosylation of Phytocannabinoids and Endocannabinoids with Os03g0702000p

Glycosylation of selected phytocannabinoids and endocannabinoids by Os03g0702000p was performed on a 200 μl scale as proof of concept. Os03g0702000p adds secondary glucose residues via β-2-1 connectivity from the primary glucose moiety; therefore, UGT76G1 was included in the reaction mixture to establish the primary glycosylation. The reactions were performed as follows. First, 1 μg of the selected phytocannabinoid or endocannabinoid substrate (1 μl of a 1 mg/ml solution dissolved in methanol, 0.016 mM final concentration) was added to 200 μl of a reaction mixture containing 50 mM potassium phosphate buffer (pH 7.2), 3 mM magnesium chloride (MgCl_2_), 2.5 mM uridine diphosphate glucose (UDPG) as the sugar donor, 10 μl of a 50% UGT76G1 glycerol stock solution (1.5 mg/ml stock solution, 1.44 μM final protein concentration), and 10 μl of a 50% Os03g0702000p glycerol stock solution (1.0 mg/ml stock solution, 0.962 μM final protein concentration). The reaction mixtures were incubated at 28 °C while shaking at 180 rpm for 18 hrs. To stop the reaction, the reaction mixture was extracted three times with 200 μl of ethyl acetate each time. The organic layers containing cannabinoid glycosides and any unreacted substrate were combined and the ethyl acetate was removed under vacuum; then, the samples were redissolved in 100 μl of 50% MeOH by volume. The reaction products were analyzed by RP-HPLC on a Dionex 3000 LC system using a Phenomenex Kinetex 5 μm XB-C18 100Å column (150 x 4.6 mm). The conditions were as follows: injection volume 50 μl, column temperature 30 °C, autosampler 8 °C, flow 1 ml/min, running time 25 min, solvent A deionized water (di-H_2_O), solvent B acetonitrile (ACN), gradient elution 10-99% ACN.

### Purification of CBD and THC Glycosides

CBD and THC glycosides were purified by C18 solid phase extraction. Hypersep C18 columns (Thermo) were prepared by hydration in methanol (MeOH), followed by 50% MeOH, then rinsed with di-H2O. UGT76G1 was deactivated and precipitated from the glycosylation reaction mixture by treatment at 95 °C for 10 minutes; then, the clarified reaction mixture was passed through the column. The column was washed successively with 10 bed volumes of di-H2O,10% MeOH, and 30% MeOH. CBD Glycoside products were eluted with 60% MeOH by volume, whereas THC glycosides were eluted with 80% MeOH by volume. The purity of the glycoside mixtures was analyzed by RP-HPLC as described above.

### Spectroscopic Analysis of Purified CBD Glycoside Mixture

CBD glycoside products produced in the glycosylation reaction with UGT76G1 were analyzed by liquid chromatography-electrospray ionization low-resolution mass spectrometry (LC/ESI-LRMS). The analysis was performed at Emery Pharma on a Shimadzu LC-MS 2010 EV instrument. The LC separation was performed on a 5 μm Silia Chrom XDB C18 150Å column (50 x 4.6 mm). The running time was 12 min and a 5-95% ACN elution gradient was utilized. LRMS was performed in positive mode with formic acid as a sample additive. The injection volume was 20 μl. Masses corresponding to the CBD aglycone up to the addition of 4 glucose residues were identified as follows:

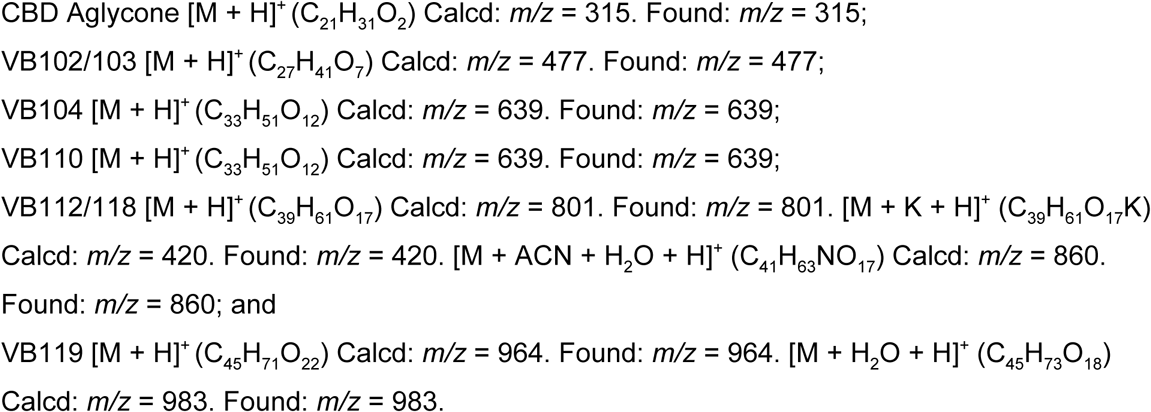

### Spectroscopic Analysis of Purified and Separated CBD Glycoside VB104

^1^H-NMR was obtained on a Bruker Avance II 400 MHz instrument using TopSpin acquisition and SpinWorks 4 processing software. 1H NMR (CD_3_OD): δ 0.81 (t, 3H, H_5”_), 1.19 (m, 4H, H_3”_, H_4”_), 1.46 (t, 2H, H_2”_), 1.54 (s, 3H, H_10_), 1.59 (s, 3H, H_7_), 1.65 (m, 2H, H_5_), 1.89 (d, 1H, H_4_), 2.41 (t, 2H, H_1”_), 2.95 (m, 1H, H_6_), 3.18-3.70 (m, 12H, H_glu*_), 3.78 (dd, 2H, H_1_), 3.94 (d, 1H, H_2_), 4.31 (m, 1H, H_9cis_), 4.42 (d, 1H, H_9trans_), 4.50 (d, 1H, H_β-1*_), 5.18 (s, 1H, H_α-1*_), 6.19 (s, 1H, H_5’_), 6.36 (s, 1H, H_3’_).

### Spectroscopic Analysis of Purified and Separated CBD Glycoside VB110

^1^H-NMR was obtained on a Bruker Avance II 400 MHz instrument using TopSpin acquisition and SpinWorks 4 processing software. 1H NMR (CD_3_OD): 0.79 (t, 3H, H_5”_), 1.16 (m, 5H, H_3_”, H_4_”), 1.47 (m, 3H, H_2_”, H_10_), 1.59 (s, 1H, H_7_), 1.65 (m, 1H, H_5_), 1.91(m, 1H, H_5_), 2.43 (t, 1H, H_1”_), 2.99 (m, 1H, H_6_), 3.16-3.67 (m, 11H, H_glu*_), 3.77 (dd, 1H, H_1_), 4.07 (m, 1H, H_2_), 4.29 (m, 1H, H_9cis_), 4.45 (d, 1H, H_9trans_), 4.49 (d, 1H, H_β-1*_), 5.23 (s, 1H, H_α-1*_), 6.59 (d, 1H, H_3’_, H_5’_).

### Spectroscopic Analysis of Purified THC Glycoside Mixture

THC glycoside products produced in the glycosylation reaction with UGT76G1 were analyzed by liquid chromatography-electrospray ionization low-resolution mass spectrometry (LC/ESI-LRMS). The analysis was performed at Emery Pharma on a Shimadzu LC-MS 2010 EV instrument. The LC separation was performed on a 5 μm Silia Chrom XDB C18 150Å column (50 x 4.6 mm). The running time was 12 min and a 5-95% ACN elution gradient was utilized. LRMS was performedin positive mode with formic acid as a sample additive. The injection volume was 20 μl. Masses corresponding with the THC aglycone along with glycosides conjugated with 2 and 3 glucose residues were identified as follows:

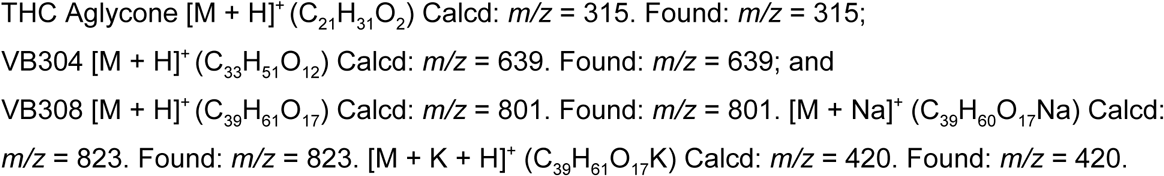

### cLogP Calculations

cLogP (logarithm of the partition coefficient between n-octanol and water) calculations were performed using ChemDraw Ultra (CambridgeSoft Corp., Cambridge, MA).

### Empirical Determination of Solubility and Detersiveness

5 milligrams of cannabosides and control chemicals were weighed out and added to 2.0ml glass HPLC vials. 500 microliters of sterile distilled water was added and the resulting solution was mixed by pipetting and gentle swirling. Solutions were then vortexed using a laboratory vortexer for 1 minute (VWR Vortex Genie, setting 10). Images were taken at the indicated steps and cropped using GNU Image Manipulation Program, and assembled using Inkscape.

---

**Supplemental Figure 1:**
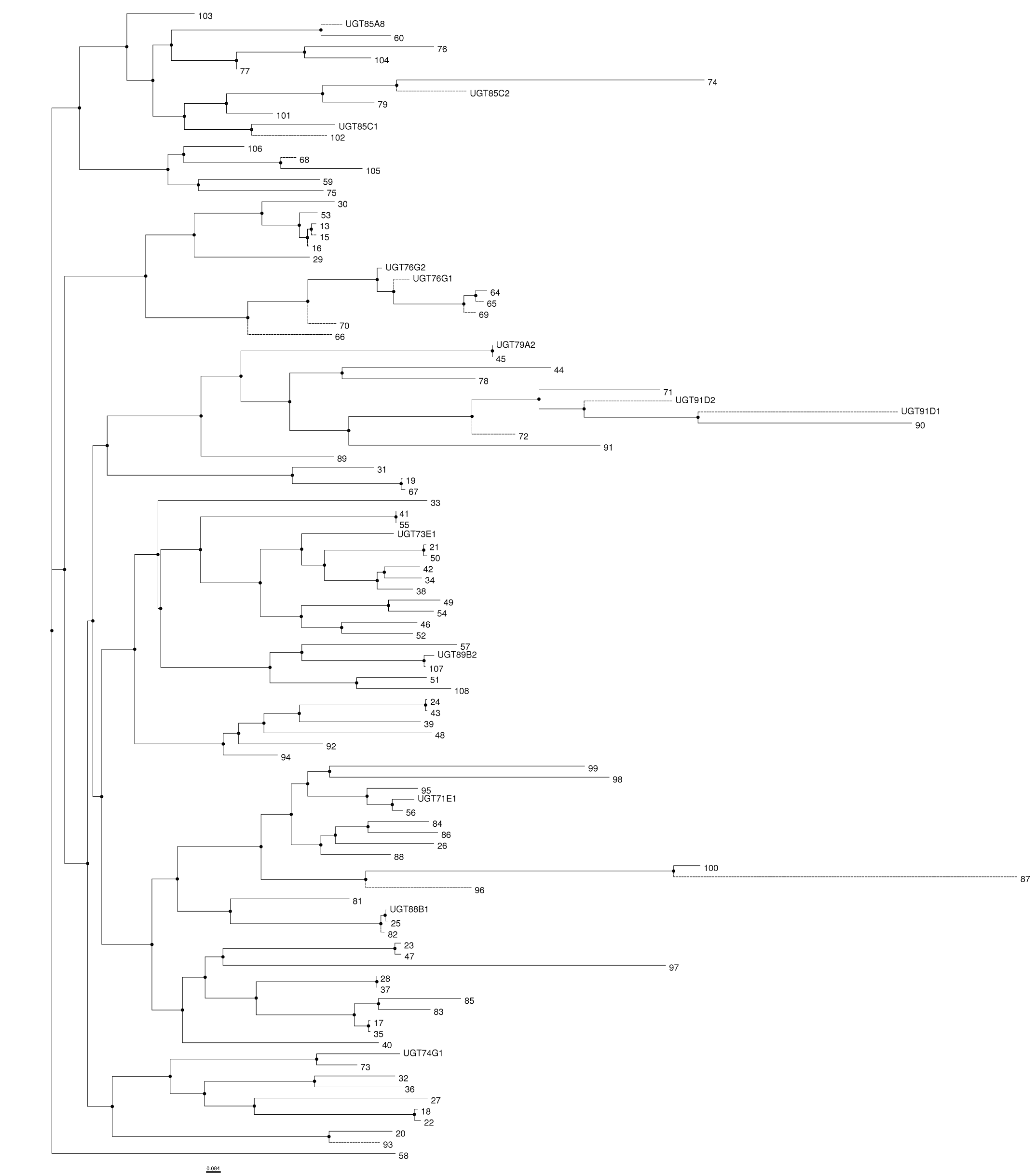
Phylogenetic tree of UGTs from Stevia rebaudiana. UGT protein sequences were mined from a Stevia transcriptome and aligned using MUSCLE in UGENE (v1.9.8). The phylogenetic tree was assembled using PHYLIP neighbor joining with Jones-Taylor-Thornton distance matrix model. Individual distances were omitted for figure clarity.

**Supplemental Figure 2:**
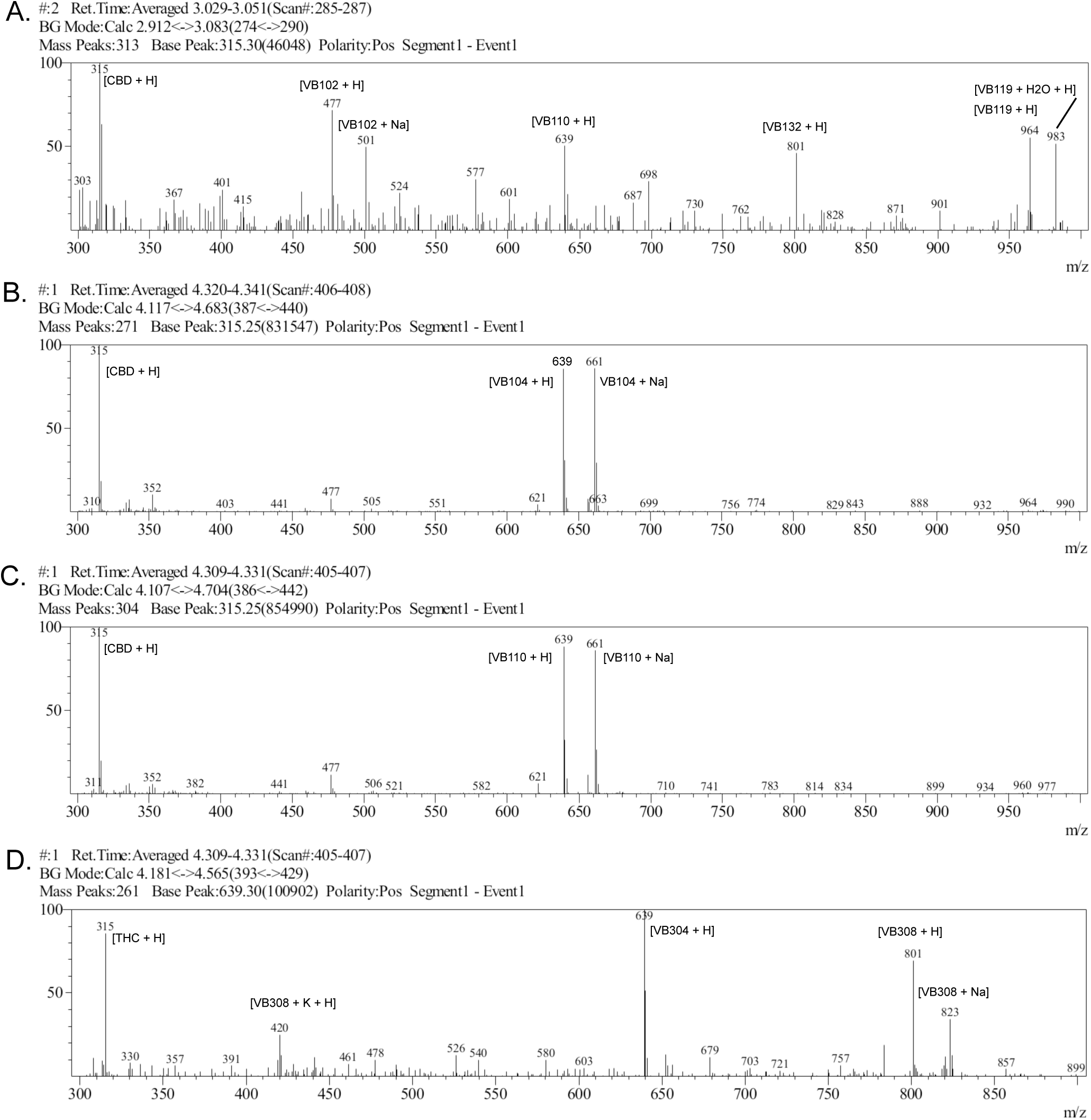
LC-ESI-MS for cannabosides. Figure 2. Mass spectra of cannabinoid glycosides generated by incubation with UGT76G1 and UDPG, where gX represents glucose plus the number of sugars attached to the parent compound. (A) Purified CBD glycoside mixture. (B) Separated CBD glycoside VB104. (C) Separated CBD glycoside VB110. (D) Purified THC glycoside mixture.

**Supplemental Figure 3:**
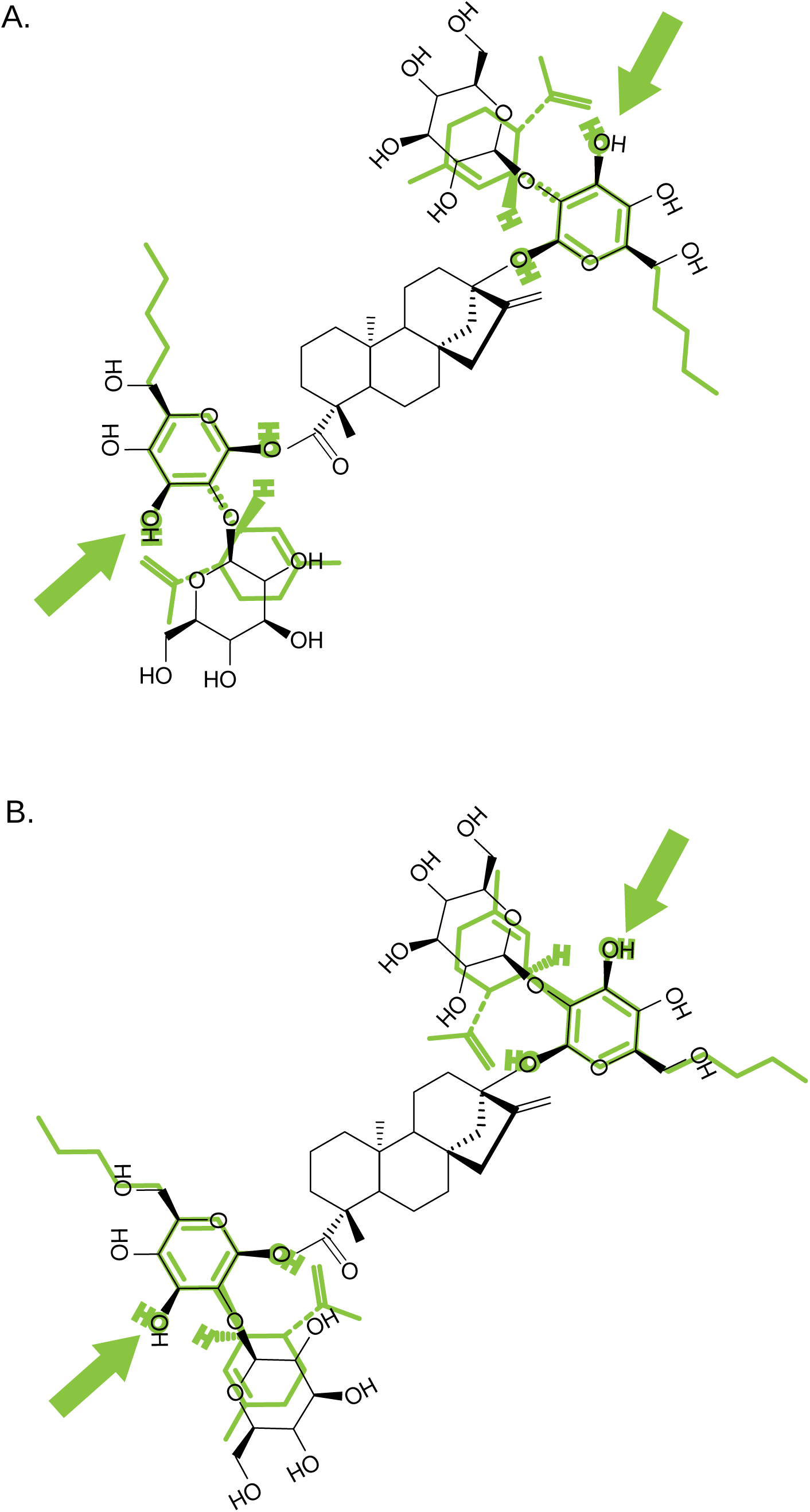
Molecular superpositioning of UGT76G1 substrates. A prototypical substrate of UGT76G1 from the stevia biosynthetic pathway, Rebaudioside E (RebE, in black), was overlaid with CBD (green) in a manner that would facilitate glycosylation of the 6'-OH group (A). RebE (black) was overlaid with CBD (green) in a manner that would facilitate glycosylation of the 2'-OH group (B).

**Supplemental Figure 4:**
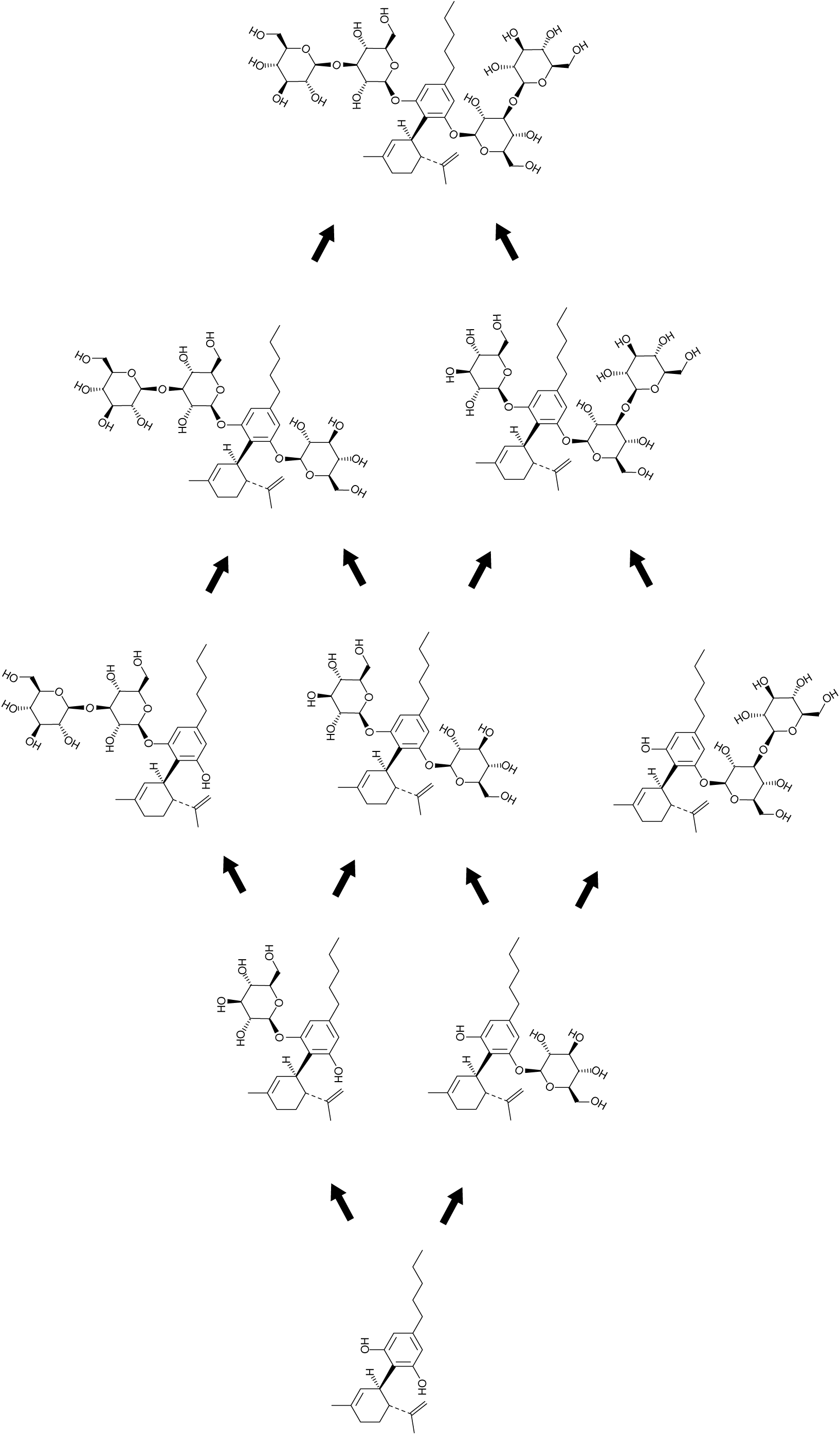
Direct glycosylation model of CBD by UGT76G1. A direct glycosylation model for the biosynthesis of CBD-glycosides by UGT76G1. CBD retains a rigid conformation and does not rotate around the C1’ axis. Glycosylations are established as the overall substrate re-positions the recipient hydroxyls towards the enzyme catalytic site.

**Supplemental Figure 5:**
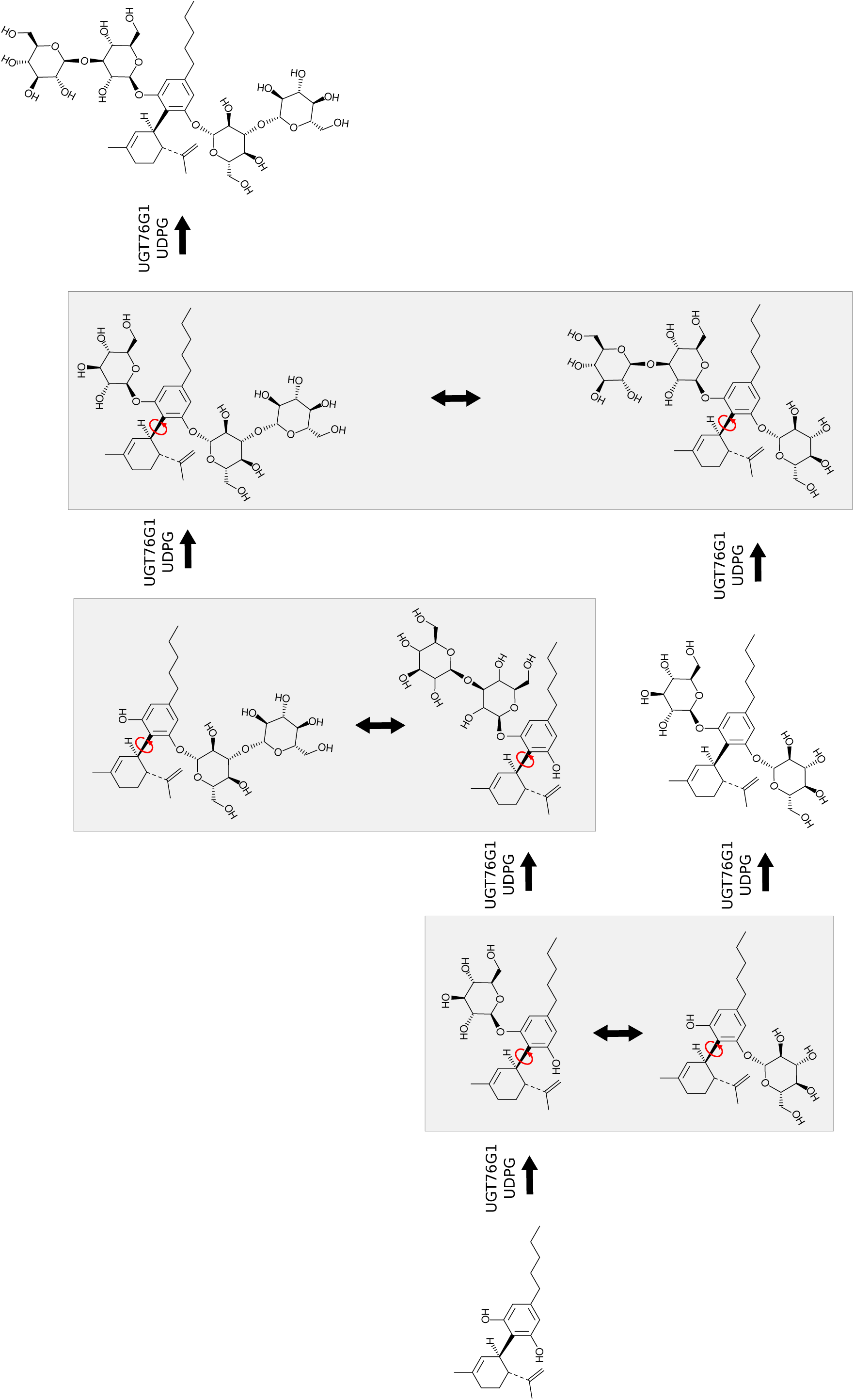
Rotational glycosylation model of CBD by UGT76G1. A rotational model for the biosynthesis of CBD-glycosides by UGT76G1. After establishing one or two glycosylations on one of the hydroxyl groups of CBD, the rotational freedom along the C1’ axis allows the resorcinol ring to rotate and swing the second hydroxyl group towards the catalytic site of UGT76G1.

**Supplemental Figure 6:**
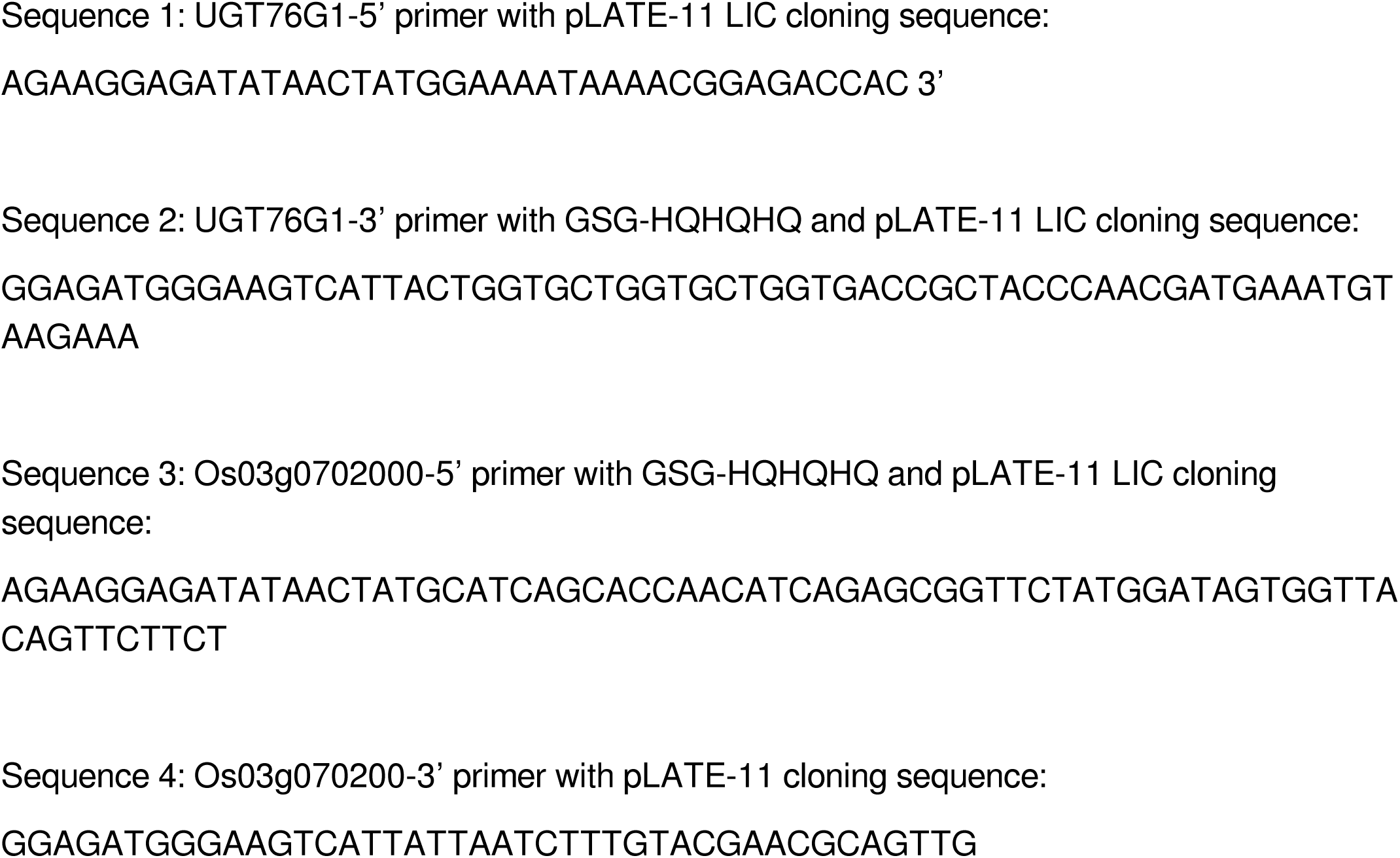
Oligonucleotide sequences. Primers used for the amplification and cloning of UGT76G1 and Os03g0702000.

